# Ocular dynamics reveal articulatory processing at single-phoneme level during silent reading

**DOI:** 10.1101/709360

**Authors:** Alan Taitz, Diego E Shalom, Marcos A Trevisan

## Abstract

Silent reading is a cognitive operation that produces verbal content with no vocal output. One relevant question is the extent to which this verbal content is processed as overt speech in the brain. To address this, we investigated the signatures of articulatory processing during reading. We acquired sound, eye trajectories and vocal gestures during the reading of consonant-consonant-vowel (CCV) pseudowords. We found that the duration of the first fixations on the CCVs during silent reading are correlated to the duration of the transitions between consonants when the CCVs are actually uttered. An articulatory model of the vocal system was implemented to show that consonantal transitions measure the articulatory effort required to produce the CCVs. These results demonstrate that silent reading is modulated by slight articulatory features such as the laryngeal abduction needed to devoice a single consonant or the reshaping of the vocal tract between successive consonants.

## Introduction

The faculty of language entails a repertoire of mental operations known as inner speech, in which verbal content is produced but voice is inhibited [1]. This internal production of words is ubiquitous: “as you read this text, the chances are you can hear your own inner voice narrating the words. You may hear your inner voice again when […] imagining how a phone conversation this afternoon will play out” [2].

One relevant question regarding speech-related tasks is the extent to which they are processed as actual speech in the brain. During speech perception, for example, neural patterns are organized around acoustic features and do not contain articulatory representations as the ones produced during speech [3]. On the contrary, speech imagery has been recently associated with the production of efference copies [2], a specific signature of motor patterns. Moreover, spectrotemporal features of inner speech were decoded with significant predictive accuracy from models built from overt speech data [4].

Increasing evidence suggests that, despite the absence of vocal articulation, motor patterns form part of inner speech. However, investigation on this matter has remained challenging, due in part to the lack of behavioral output of speech imagery, which makes difficult to time-lock precise events (acoustic features, phonemes, words) to neural activity [4]. Silent reading offers a direct way to tackle this problem, since it provides us with the trajectory of the reader’s eyes along the text as a natural behavioral output [5]. Although a fair amount of behavioral experiments demonstrated that phonology affects silent reading [6,7], a quantitative study on the relation between articulatory and ocular variables is still lacking.

Here we hypothesize that articulatory simulations at single-phoneme level underlie silent reading. We tested this by a quantitative exploration of ocular and articulatory dynamics during the reading of consonant-consonant-vowel (CCV) speech structures.

Ocular trajectories basically consist of gaze fixations separated by rapid eye movements called saccades [8]. During reading, each word usually receives a few fixations, between which neighboring words may be also fixated [9]. Here we characterized reading dynamics by computing standard timing variables such as the duration of first fixations and the integration of the successive fixations on a word, which have been largely used to disentangle visual, lexical and contextual processing in reading [10].

To emphasize articulatory dynamics, focus was put on CCVs formed with plosive consonants, which require sharp movements to close the vocal tract completely. For instance, when the lips occlude the air passage, consonant *b* is produced; in this case, also the vocal folds are active. Instead, when folds are inactive, the same occlusion produce the unvoiced plosive *p* [11]. Plosives therefore involve sharp articulatory movements combining on-off folds activities [12], and sequences of plosives and consonants have also been modeled and synthesized from physical principles [13,14]. From a lexical point of view, plosive CCVs form a homogeneous subset of pseudo-words with low intra-word frequency values [15]. This combination of motor, mathematical and lexical features makes these CCVs an ideal set of stimuli to explore the relationship between silent reading and vocal articulation.

## Results

Thirty native Spanish speakers read a set of consonant-consonant-vowel structures (CCV) from a computer screen. The set was formed by every combination of fricative consonants (*s*, *f*, *j*) and plosive consonants (*b*, *d*, *g*, *p*, *t*, *k*) ending with the vowel *a*. A few examples of these structures are *fga* (fricative-plosive), *bda* (plosive-plosive) and *jsa* (fricative-fricative), shown in Figure 1a. Intra-word frequency of each structure are displayed in Figure 1b (none is a Spanish word).

**Figure 1.**
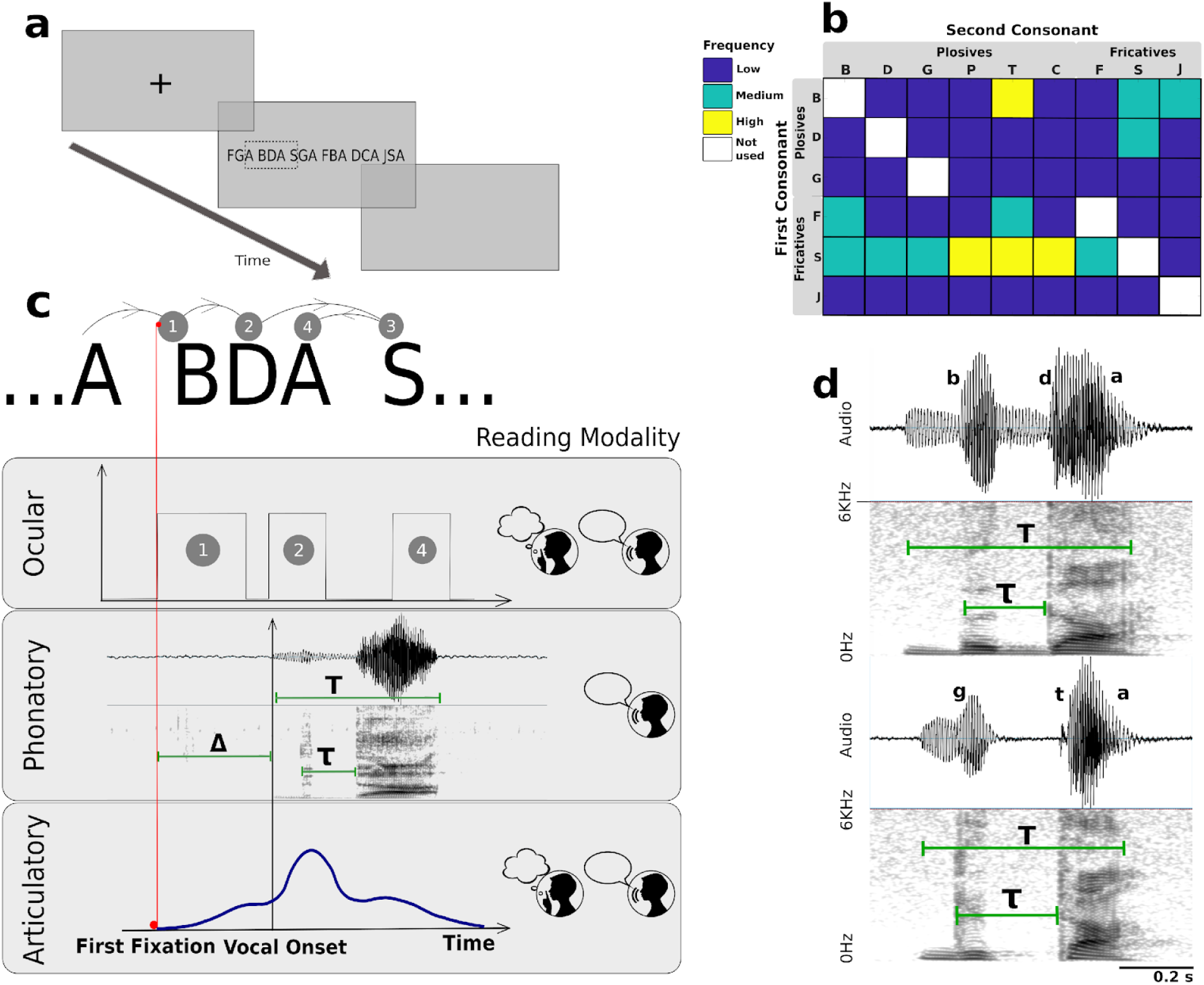
Monitoring ocular, phonatory and articulatory variables during reading. **a.** After fixing the sight on a cross in the middle of the screen, a sequence of 6 CCVs appeared for the participant to read. The operation was repeated until three repetitions of the complete CCV set was presented. In one block, participants read aloud and in the other block they read silently. **b**. Intra-word frequency *f* of the CCVs in Spanish (in appearances per million words). The range was discretized in three levels: low (*f*<10), medium (10<*f*<50) and high (*f*>50). **c**. Three types of variables were measured during reading. Ocular: duration of first fixation (FFD=1); duration of fixations on a CCV prior to fixating a neighboring one (FPRT=1+2); and total fixation duration (TFT=1+2+4). Phonatory: We measured the delay between the first fixation and the vocal Onset Δ, the transition time between consonants τ, and the total pronunciation time *T*(overt reading). Articulatory: Lip movement was registered by an accelerometer fixed to the lower lip. **d**. Spectrograms and phonatory variables of *bda* (voiced plosives) and *gta* (unvoiced second consonant).

The experiment was divided into two blocks, each formed with three repetitions of the complete set of CCVs. In one block the participants read the set aloud, and in the other they were instructed to read in silence. Blocks and CCVs were randomized before presentation. For each CCV we measured the following variables, sketched in Figure 1c (videos available at Supplementary Information):

<u>Ocular</u> (aloud and silent blocks). Eye movements were recorded and the following variables were computed for each CCV: 1. the duration of the first fixation (FFD); 2. the duration of all fixations prior to passing to another CCV (First Pass Reading Time, FPRT) and 3. the total fixation time (TFT). These are standard ocular variables used to characterize cognitive processing in reading; the first two are typically considered early measures, whereas measure 3 reflects later processing stages [16,17].
<u>Phonatory</u> (aloud block). From the spectrograms of the audio records, we computed: 1. the delay Δ between the onset of visual fixation and voice onset; 2. the total duration *T* of the CCV, and 3. the consonantal transition τ, shown in detail in Figure 1d for the plosive-plosive combinations *bda* and *gta*. We defined τ as the interval between the release of the first occlusion, characterized by a brief voiced sound, and the release of the second one, that give rise to the vowel *a*. Finally, differences in speech rate across subjects were washed out by normalizing the transitions to the total duration of the CCV, τ ′ = τ/*T*, leaving us with the pre-phonatory variable Δ and the phonatory variable τ′.
<u>Articulatory</u> (aloud and silent blocks). Lip movement was acquired by attaching an accelerometer to the lower lips with medical tape.

## Articulation is inhibited during silent reading

We first investigated the articulatory activity in both aloud and silent reading blocks. The main vocal tract articulators are the lips, the jaw and the tongue. Lip movements can be easily obtained from the exterior of the mouth, minimizing the interference of the measuring device with articulatory dynamics. Since articulators are coordinated during speech [18], we assumed that the range of lip movement is a good estimate for the range of the whole articulatory movements.

In Figure 2 we show the dynamics of the lips for the CCVs that start with the bilabial consonant *p* (*pba*, *pda*, *pga*, *pta* and *pca*), which requires maximal lip displacement. In Figure 2a we show the absolute value of the accelerometer signals during reading aloud, aligned to the beginning of the vocalization. We recovered the characteristic dynamics of the lips, with movement starting roughly half a second before phonation, and maximum displacement at phonation onset [19]. The fact that maxima mostly occur at vocal onset can be observed also in Figure 2b, where the signals were aligned to their maximum value instead of the vocal onset. We take advantage of this to compare the signals of the aloud block (red) with those of silent block (blue), for which we do not have the vocal onset reference. Two groups are readily identified among the silent readers (blue), one with low and the other with virtually null lips activity. Direct comparison between modalities shows that, when present, articulatory movements during silent reading are roughly two orders of magnitude smaller than those of reading aloud. These results allow us to explore how articulatory processing affects silent reading, in a context of strong inhibition of vocal articulation.

**Figure 2.**
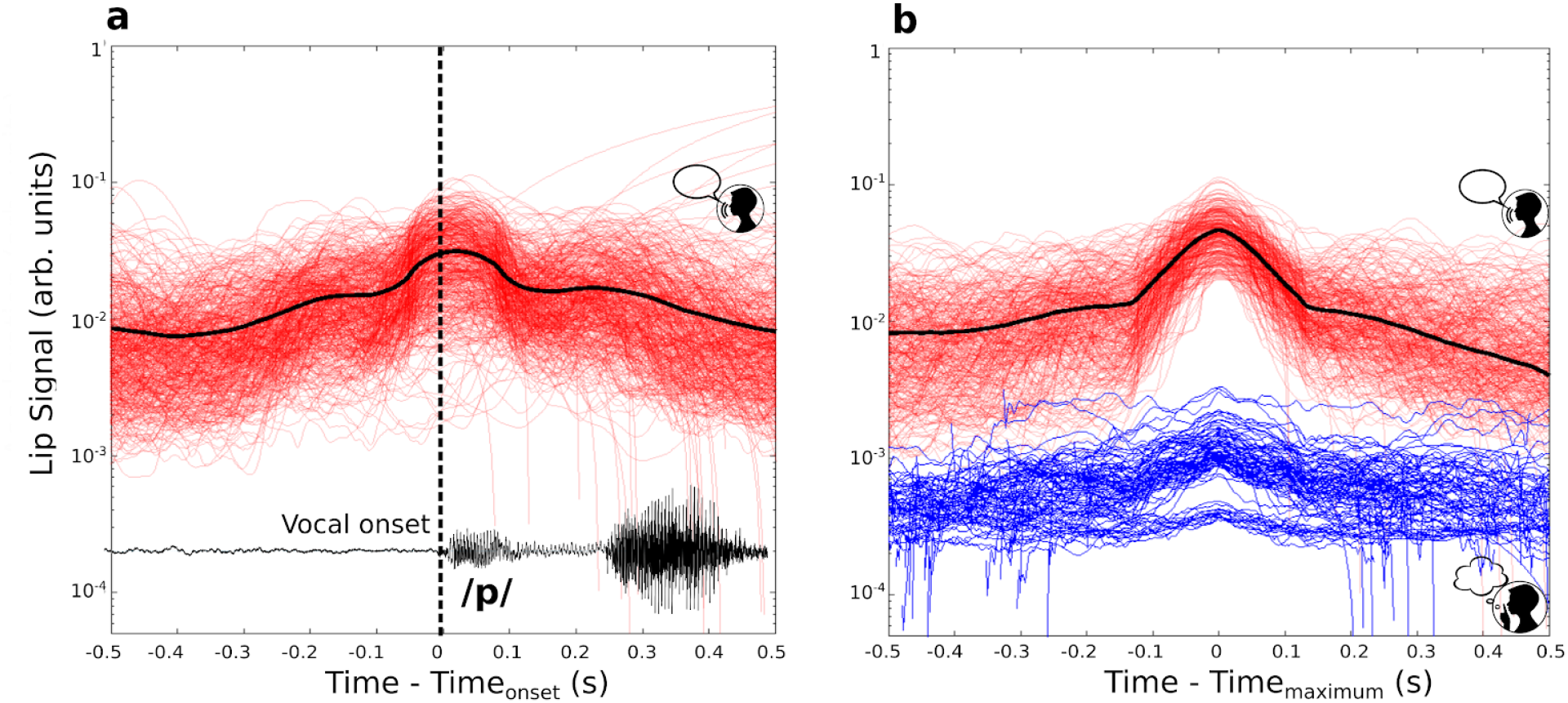
Lip movement is strongly inhibited during silent reading. **a.** Red traces are the accelerometer signals during reading aloud, aligned at vocal onset (absolute value, smoothed by a 140 ms moving window). Single trials are represented in red and averaged in black. **b.** In order to compare ranges of articulation of the different reading modalities, the signals were realigned to the maximum. Aloud reading is displayed in red and its average in black while silent reading in blue. At maxima, traces exhibit a difference of roughly two orders of magnitude.

## Silent reading is modulated by consonantal transitions

In this section we analyze the effects of phonation and memory on the ocular dynamics during reading. We focused on the CCVs formed with plosive consonants, for which articulatory movements are emphasized with respect to other consonantal families. The analyses for these other families (plosive-fricative combinations) show milder effects and can be found in Supplementary Information.

We used the consonantal transition τ′ to characterize the phonation of the CCVs; since participants read each CCV six times (three times per block), the repetition number was used to test for short term memory effects. Long term memory effects were examined using the frequency of appearance of the CCVs in a Spanish corpus. Statistical tests show no effects of frequency on any visual variable (last column of Table 1) as expected by the low frequency levels of the whole set of plosive CCVs (with the exception of BTA), as sketched in Figure 1b.

**Table 1:**
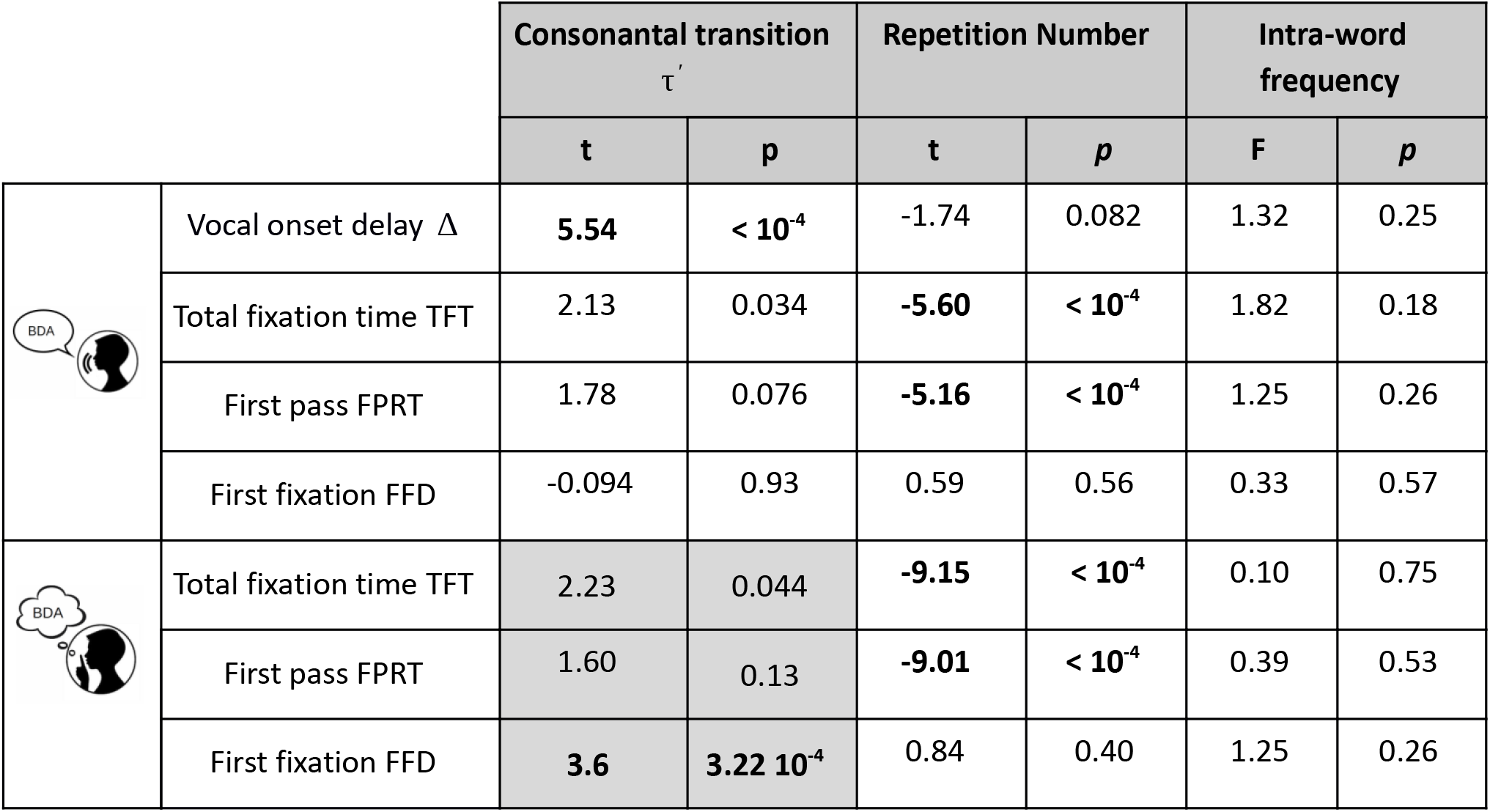
Effects of phonology and memory on ocular variables during reading. An ANOVA test revealed no significant effect of intra-word frequency on any of the variables. **Aloud block**. A multiple linear regression was conducted for the onset delay Δ and for each of the ocular measures, with consonantal transition and repetition number as independent variables. **Silent Block.** To compare variables across blocks, two different tests were performed. Short term memory effects on the ocular measures was accounted for by performing a linear regression test with repetition number as the independent variable. Ocular variables (silent block) and consonantal transitions (aloud block) were averaged across participants and repetitions before performing a linear regression test weighted by the statistical errors (gray background).

| | | Consonantal transition<br>$\tau'$ | | Repetition Number | | Intra-word frequency | |
| --- | --- | --- | --- | --- | --- | --- | --- |
|  |  | t | p | t | p | F | p |
| | Vocal onset delay $\Delta$ | 5.54 | $< 10^{-4}$ | -1.74 | 0.082 | 1.32 | 0.25 |
| | Total fixation time TFT | 2.13 | 0.034 | -5.60 | $< 10^{-4}$ | 1.82 | 0.18 |
| | First pass FPRT | 1.78 | 0.076 | -5.16 | $< 10^{-4}$ | 1.25 | 0.26 |
|  | First fixation FFD | -0.094 | 0.93 | 0.59 | 0.56 | 0.33 | 0.57 |
| | Total fixation time TFT | 2.23 | 0.044 | -9.15 | $< 10^{-4}$ | 0.10 | 0.75 |
| | First pass FPRT | 1.60 | 0.13 | -9.01 | $< 10^{-4}$ | 0.39 | 0.53 |
| | First fixation FFD | 3.6 | $3.22 \cdot 10^{-4}$ | 0.84 | 0.40 | 1.25 | 0.26 |

### Aloud block

We analyzed the effects of phonation and memory on the ocular variables and also on the pre-phonatory variable Δ during reading aloud (Table 1, top row). For this last case, we found a significant effect of τ′ on Δ, as reported in a previous work [15]. This means that the preparatory stages before phonation are longer for CCVs requiring longer consonantal transitions, which has been identified as a possible effect of articulatory processing [15]. For the ocular variables, linear regressions were conducted using consonantal transition τ′ and repetition number as independent variables. A negative dependence with repetition number emerged on FPRT and TFT, and no effects were found on FFD, supporting that first fixation durations are not sensitive to habituation during reading aloud. Finally, we found no effects of transitions τ′ on the ocular variables. This result was expected, given the complexity of ocular dynamics during reading aloud, in which the eyes appear to be holding in place for many fixations “so as to not get too far ahead of the voice” [20].

### Aloud block vs. silent block

We next concentrated in the relationship between silent and overt reading, which is the main aim of this work. For this sake, we compared ocular data from the silent block with phonological data from the aloud block (Table 1). To compare cross-block variables, we first performed linear regressions for each ocular measure using repetition number as the independent variable. Consistent with the results obtained for the aloud block, this revealed that FPRT and TFT systematically decrease across repetitions, while FFD presents no short term memory effects. This gives us confidence that the variables that integrate the fixations on a CCV (FPRT and TFT) are affected by habituation while the duration of the first fixation FFD is not, making the latter a good candidate for articulatory processing. To test this, we compared the ocular variables of the silent block with the phonatory variable τ′ of the aloud block, averaging across participants and repetitions. This is summarized in Figure 3, where we show each ocular variable as a function of consonantal transitions per CCV. A strong positive relation emerged for FFD (Figure 3a), while for TFT and FPRT the relation did not reach significance (Figure 3b and c).

**Figure 3.**
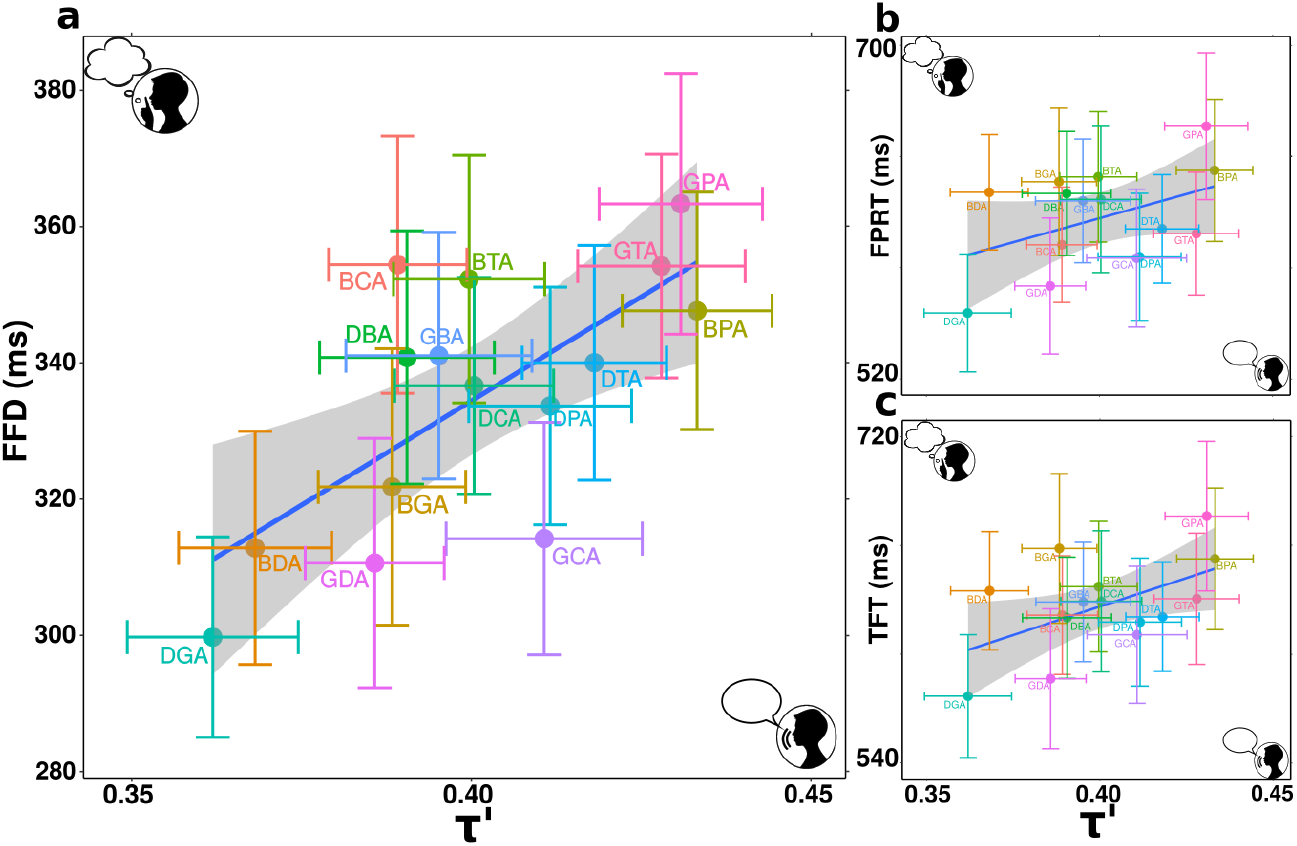
Silent reading is modulated by consonantal transitions. **a.** First fixation durations from silent block as a function of transition τ′ from aloud block. Linear regression was conducted using averaged CCVs points with its respective statistical standard error considered, showing that strong dependence between FFD and τ′ (*t* = 3.60, *p* = 3.2210^−3^) **b-c.** First pass reading time (FPRT) and total fixation time (TFT) as a function of τ′, FPRT: (*t* = 1.60, *p* = 0.13); TFT: (*t* = 2.30, *p* = 0.044).

Taken together, these analyses reveal that the duration of the first fixations FFD in silent reading are not affected by habituation, and are tightly correlated with the consonantal transitions τ′. This supports that ocular dynamics during silent reading are strongly modulated by phonatory features produced during the actual utterance of the speech structures.

## Consonantal transitions measure articulatory efforts

So far, our results support that silent reading is strongly modulated by the transitions τ′ between consonants. Here we connect this timing variable with the motor actions needed to produce the utterance of a CCV. Like any other speech structure, producing a CCV requires two basic motor actions: the larynx needs to be abducted to control the oscillations of the vocal folds, and the vocal tract shape is reconfigured to produce the different phonemes. We explored how the consonantal transitions τ′ are related to these specific motor actions.

### Laryngeal effort

The vocal folds are a pair of elastic membranes located at the glottis, within the larynx. The folds can be set into oscillatory motion by the passing airflow, raising the lung pressure over a threshold that increases with glottal abduction. Within our set of CCVs, vibration of the folds is either active during the whole vocalization (*bda*, *bga*, *dba*, *dga*, *gba*, *gda*) or inactivated during the second plosive (*bca*, *bta*, *bpa*, *dca*, *dta*, *dpa*, *gca*, *gta*, *gpa*), as evidenced by the traces of the fundamental frequency in the spectrograms of Figure 1d. To produce the CCVs in the second group, the glottis needs to be actively abducted to prevent vibration during the middle unvoiced plosive, which involves a devoicing effort. Consonantal transitions reflect this, with a smaller mean duration for the former group, τ′ = (38.2 ± 0.3) 10^−2^ than for the latter, τ′ = (41.3 ± 0.3) 10^−2^ (t-test: *t* = −4.99, *p* < 10^−4^).

### Tract effort

A CCV formed by plosives is produced by two successive occlusions that occur while the tract evolves towards the vowel *a*. This produces slight differences in the articulatory effort when consonants are interchanged. For instance, *bda* requires less effort than *dba* because in the latter, the lips’ closure for the *b* occurs closer to the vowel *a*, for which the mouth is fully opened, requiring a larger vocal tract deformation. These slight vocal tract asymmetries are also reflected by consonantal transitions, with τ′ = (37.3 ± 0.7) 10^−2^ for the group (*bda*, *bga*, *dga*) and τ′ = (39.0 ± 0.7) 10^−2^ for the group with interchanged consonants (*dba*, *gba, gda*), with a trend towards significance (*t* = 1.79, *p* = 0.074). Beyond this subtlety, the main variations in the tract effort arise from the anatomical differences between plosives: for instance, producing a *g* requires displacing the body of the tongue, while for a *d* only the tip is shifted. These anatomical differences can be accounted for using mathematical functions *A* (*x*, *t*) for the cross section of the vocal tract along its length *x* from the entrance to the mouth.

In a previous work [15] we capitalized on this mathematical description, joining together the laryngeal and vocal tract components in a single equation for the vocal effort *E* that reads:

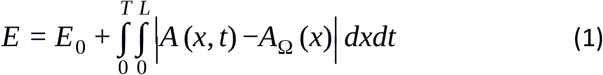

The first term refers to the laryngeal component that takes a value *E*_0_ > 0 when a phoneme is devoiced, and *E*_0_ = 0 otherwise. The second term accounts for the elastic forces produced by deformations of the vocal tract, from a neutral shape *A*_Ω_ (*x*) to a general shape *A* (*x*), as sketched in Figure 4a. The tract effort is obtained by integrating the section *A* (*x*, *t*) along the duration *T* of a CCV, from the vocal tract entrance at *x* = 0 to mouth exit at *x* = *L*. In this way, equation 1 provides an estimate of the total laryngeal and vocal tract articulatory actions needed to produce our speech structures.

**Figure 4.**
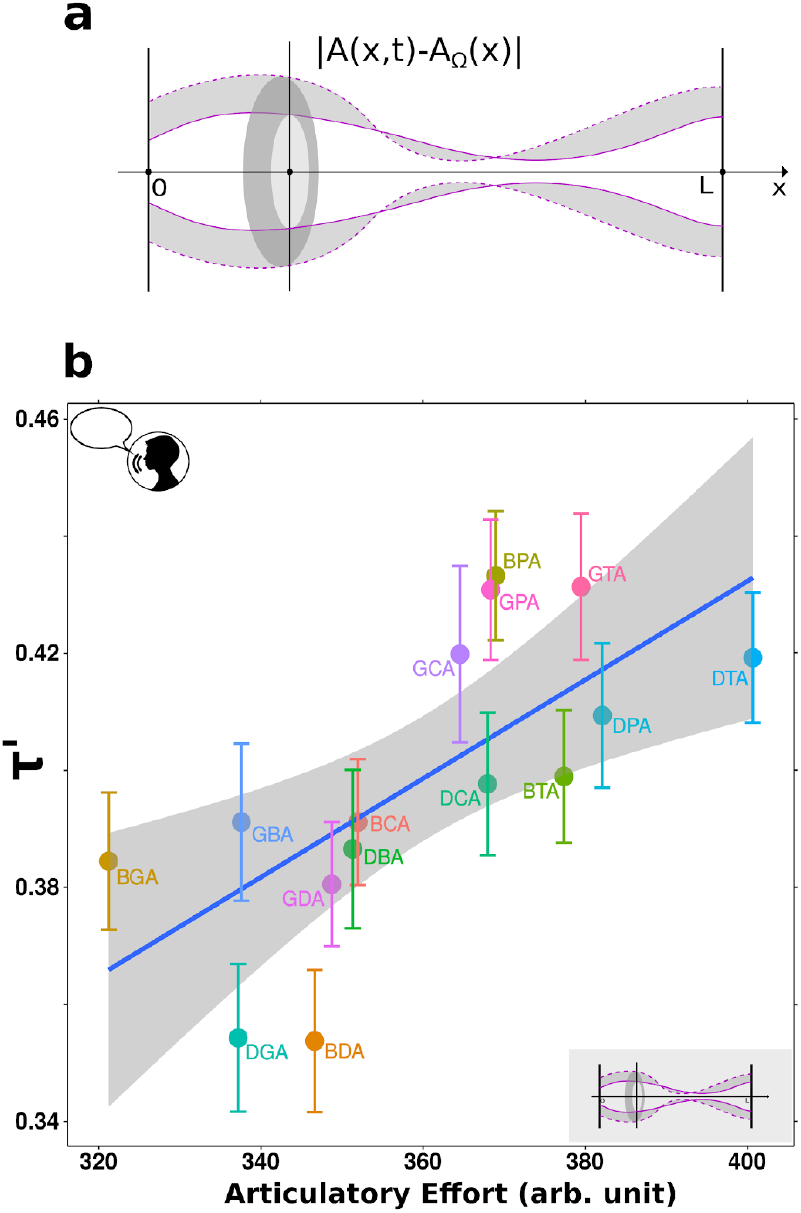
Computing total vocal effort from consonantal transitions. **a.** The shape of the vocal tract is described by its cross section *A* (*x*, *t*), where *x* is the distance from the vocal tract entrance (*x* = 0) to the exit of the mouth (*x* = *L*). The articulatory effort needed to pronounce a CCV can be estimated from the deformations of the vocal tract (shadowed region) with respect to a relaxed configuration (dotted line). **b.** The transitions τ′ measured on the recorded CCVs averaged across participants, exhibit a positive relation with the modeled articulatory effort computed with Equation 1 (*t* = 3.4, *p* = 4.7 10^−3^).

To compute the efforts of each CCV, we used vocal tract functions *A* (*x*, *t*) and devoicing efforts *E*_0_ from anatomical data as reported elsewhere [15,21,22]. In Figure 4b we show the estimated efforts *E* for each CCV as a function of τ′ (averaged over participants and repetitions). This result supports that consonantal transitions are a good estimate for the vocal effort needed to produce a CCV at laryngeal and vocal tract levels.

Taking together the analyses of the previous sections, our results show that silent reading involves the processing of precise articulatory features. First fixations during silent reading are lengthened when the CCVs require a greater laryngeal and/or articulatory effort to be pronounced, even when this implies devoicing a single consonant or changing the point of occlusion in the vocal tract.

## Conclusions and Discussion

The faculty of language involves the ability to switch between inner and actual speech, offering a unique opportunity to study the differences in motor processing between these operations. We capitalized on this faculty to explore the articulatory signatures during aloud and silent reading. Our main findings are that 1. the duration of the first fixations on the CCVs in silent reading are strongly correlated with the duration of the consonantal transitions when the CCVs are read aloud, and 2. consonantal transitions measure the vocal effort needed to pronounce the CCVs. Together, these results support that short pseudowords are processed at single-phoneme articulatory level during silent reading.

Two theories have accounted very differently on the role of the motor system during inner speech tasks. The abstraction theory supports that inner speech only activates abstract linguistic representations, independently from any articulatory mental simulation [23,24]. On the other hand, the motor simulation theory [25,26] describes inner speech as an activity that involves a similar motor processing than overt speech, including articulatory detail. Halfway between both theories, Oppenheim and Dell proposed a flexible abstraction hypothesis, in which inner speech operates at two levels: an abstract processing level, and one that incorporates a lower-level articulatory planning [27].

Recent investigations accumulate evidence towards motor simulations; Whitford et al [2] showed that speech imagery is associated with an efference copy with detailed auditory properties, treating it as a kind of action. In the same direction, a groundbreaking result was reported by Anumanchipalli and colleagues [28], who mounted a neural decoder based on articulatory dynamics capable of synthesizing speech when a participant silently mimed sentences. Investigation on inner speech has direct implications in the development of devices to restore speech [29], and also in other fields as psychiatry, where failure in the monitoring of inner speech has been associated with verbal hallucinations in schizophrenia [30].

However, not so much attention has received the motor processing during silent reading. Previous research showed that silent reading activates phonological representations in the brain [6], and also that global aspects of overt speech affects the dynamics of silent reading [7]. In line with this research, our work is a quantitative in-depth study of the articulatory signatures during silent reading. We have shown that slight motor variations are sufficient to affect the reading dynamics and indicate that this task, as speech imagery, is assisted by precise vocal simulations.

## Materials and Methods

### Participants

Thirty native Spanish speakers (13 females and 17 males, age range 22–33, mean age 27), undergraduate and graduate students at the University of Buenos Aires with normal vision, no speech impairments and fluent reading skills completed the experiment. Participants signed a written consent form. All the experiments described in this paper were approved by the CEPI Ethics Committee of the Hospital Italiano de Buenos Aires, qualified by ICH (FDA-USA, European Community, Japan) IRb00003580, and all methods were performed in accordance with the relevant guidelines and regulations.

### Stimuli and tasks

The experiment consisted of two blocks separated by a three minute break, each involving a single reading modality. Before the aloud reading block, participants were instructed to say the pseudo-words as they appeared on the screen. Before the silent reading block, participants were instructed to read the sequences silently, as they normally do when reading a book. The order of the blocks was randomized.

The experimental design was identical for each block: a screen containing a sequence of 6 randomized consonant-consonant-vowel (CCV) structures was presented right after the participant fixated a red point (Figure 1a). Once the CCVs were read, the participant pressed a key and the process restarted until completing the set (videos are available as Supplementary Materials).

The CCVs were built using fricative and plosive consonants followed by the vowel *a*. We used the most common Spanish fricatives *f*, *s* and *j* (pronounced as **f**ace, **s**tand, and the scottish lo**ch**) and all the Spanish voiced plosives *b*, *d, g* (**b**ay, **d**ye and **g**ray) and unvoiced plosives *p, t, c* (**p**ay, **t**ie and **c**ray). Examples of these structures are *fsa* (fricative-fricative), *sda* (fricative-plosive) and *bta* (plosive-plosive). We excluded CCVs that repeat the same consonant, which makes a total number of combinations of 9 × 8 = 72 CCVs. Here we analyzed the 15 CCVs formed by a voiced plosive (*b*, *p*, *d*) followed by any other plosive of the set (*t*, *g*, *c*, *b*, *p*, *d*). The reason is that in these combinations, vocal folds vibrate the initial voiced plosives, allowing us to determine precisely the beginning of the CCV in the spectrogram. Unvoiced plosives, on the contrary, leave no sound or spectrogram traces and were excluded from the analysis. Voiced plosive-plosive combinations were repeated three times per block (15 CCVs 3 repetitions = 45), while all other 57 CCVs were repeated randomly two or three times-adding 135 repetitions- to make a total of 180 stimuli (6 CCVs per screen 30 trials) per block. Data was recorded for 10800 CCV samples (30 subjects 2 blocks 180 CCVs). In order to avoid screen edges effects on the ocular variables, the first and last CCVs on the screen were discarded (Figure 1a). The final database was composed by 7200 CCV samples.

The frequency *f* of each combination of consonants was computed from a large Spanish corpus [31], as the intra-word appearances per million words. Since the range of frequency values was large (0 ≤ *f* ≤ 10^5^), the range was discretized in three levels: low (*f* ≤ 10), medium (10 ≤ *f* ≤ 50) and high (*f* ≥ 50). None of the CCVs are Spanish words. Courier New monospaced font was used to ensure a fixed-width of 24 pixels for every character. Stimuli were presented in black over a white background.

### Eye tracking data

Visual trajectories were acquired through a desktop-mounted, video-based eye tracker (EyeLink II; SR Research Ltd., Kanata, Ontario, Canada) at a sampling frequency of 1 kHz in binocular mode (nominal average accuracy 0.5°, space resolution 0.01° RMS). The stimuli were presented on a 19 in. monitor model Samsung SyncMaster 997 MB, at an eye–monitor distance of 50 cm. Eye position was recorded at a resolution of 1024 × 768 px in all tasks. The head was stabilized with a forehead rest.

Eye movements during reading include saccades, which are rapid eye movements, and fixations, where the eyes remain relatively still for about 200–300 ms. In this experiment we computed three of the most frequent variables used for measuring processing time [20]: 1. First Fixation Duration (FFD), 2. First Pass Reading Time (FPRT), which is the sum of all fixation durations on the word before any other word is fixated, and 3. Total Fixation Time (TFT), the sum of all fixations durations on a word.

### Acoustic and articulatory data

Sound was recorded with a commercial microphone placed at 0.3 m from the head of the participant. Lip movement was recorded using an accelerometer ADXL335 fixed to the lower lip with medical paper tape. Recordings were made in the direction of maximum amplitude of lip movement, along the perpendicular axis of the accelerometer. Sound and articulatory signals were acquired with a DAQ (USB-1608FS-Plus, Measurement Computing, Norton, Massachusetts), which ensured synchronization of both inputs and a maximum onset delay of 50 ms. Psychtoolbox library from MATLAB was used to synchronize the eye tracker and DAQ system to the computer clock.

For each CCV, we measured the following three timing variables from the audio records:

a. The delay Δ between the first fixation and the vocalization onset. Vocal onsets were established at the appearance of the spectral signature of the initial consonant (noisy spot for fricatives and fundamental frequency for voiced plosives).
b. The duration *T* of the uttered CCV
c. The transition τ during the reconfiguration of the vocal tract from the first to the second consonant. We followed the procedure described in [15] to define the transitions τ in the time-frequency domain. For voiced plosive-plosive combinations, as shown in Figure 1d, τ goes from the release of the first plosive, characterized by a voiced sound structure, to the release of the second one into the vowel *a;* for plosive-fricative combinations, τ is the interval bounded by the brief broad-band plosive noise and the purely noisy fricative spot, as shown for *gfa* in Supplementary Figure S1a; for fricative-plosive combinations, the transition is the interval between the abrupt end of the fricative and the release of the plosive into the vowel, as shown for *fca* in Supplementary Figure S2a.

To wash out the speech rate differences between speakers, normalized transitions τ ′ = τ /*T* were used throughout this work.

From the recorded 2700 voiced plosive-plosive combinations, we discarded those that were not fixated with the eyes, the mispronounced ones and those that contained more than one attempt to produce a CCV. We have also excluded 2 participants who mispronounced more than 45% of the CCVs throughout the experiment, leaving a dataset of 2415 CCVs. We then discarded repetition which presented fixation durations (FFD, FPRT and TFT), delay times Δ or transitions τ′ out of the 95% confidence interval (<u>*x*</u> ± 2*s*) from the participant’s mean value of each variable. A final dataset of ocular, acoustic and articulatory data of 2178 CCVs from 28 participants was used to perform the analyses throughout this work.

### Vocal tract model

During a sequence of vowels and *n* plosive consonants, the vocal tract area *A* (*x*, *t*) can be mathematically described by [21]:

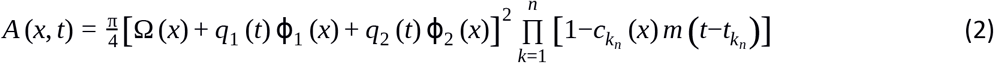

The squared factor represents the vowel substrate, with empirical functions Ω, **ϕ**_1_ and **ϕ**_2_ obtained from an orthogonal decomposition calculated from MRI anatomical data. Any sequence of vowels is generated by the evolution of *q*_1_ and *q*_2_. Plosive consonants are generated through the bell-shaped functions *c*_*k*_ and *m*. These functions reach the value 1 around *x* = *x*_*k*_ and *t* = 0 respectively, producing a specific occlusion *A* (*x*_*k*_, *t*_*k*_) = 0. The specific features of each plosive is represented by the spatial functions *c_k_* that characterize the anatomy of the occlusion and also by the time function *m* that represents the temporal dynamics of the occlusion (activation-deactivation).

## Supporting information

Example Video

## Acknowledgements

We thank Juan Kamienkowski for his time and willingness during the experiments. The research reported in this work was partially funded by the National Scientific and Technical Research Council (CONICET), the University of Buenos Aires (UBA).

