## Supplementarty Materials for "Ocular dynamics reveal articulatory processing at single-phoneme level during silent reading"

**Supplementary Material**

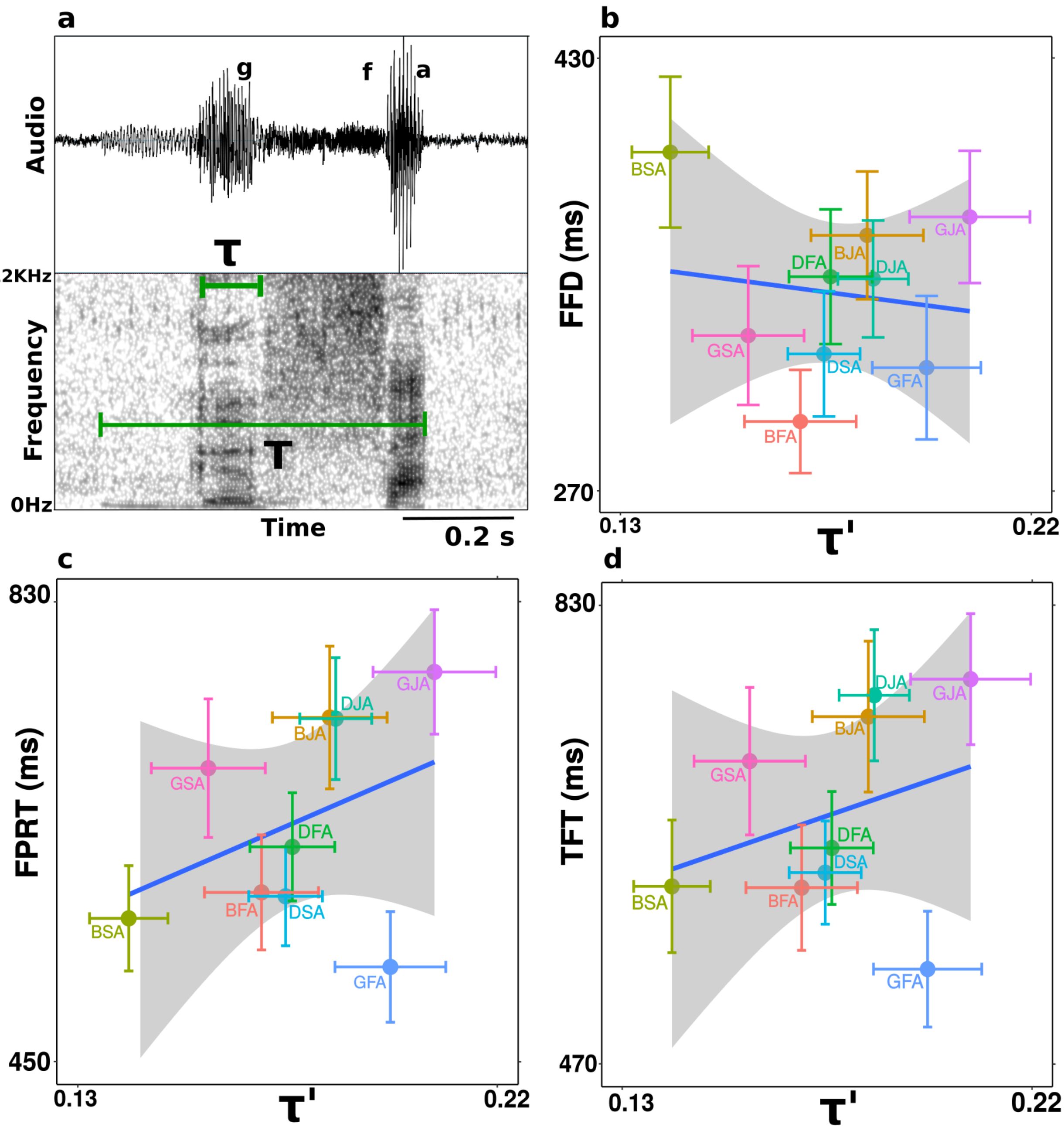


**Figure S1 | Voiced plosive-fricative combinations. a.** Audio and spectrogram example from /GFA/ pronunciation. Transitions are characterized by the interval that goes from releasing the occlusion of the plosive to the formation of a constriction to generate the fricative. Spectrally, the transition is described by the voiced sound structure produced after the plosive, and before the formation of the purely noisy fricative spot. **b - d**. Linear regression test was conducted for the variables TFT, FPRT and FFD with consonantal transition  as the independent variable. Means and standard errors per CCV were considered for the regression as data points come from different block. Although a positive tendency is observed, any of the variables hold a statistically significant relation with $\tau'$**:** FFD $(t=-0.16, p=0.87)$ FPRT $(t=1.07, p=0.32)$ and TFT $(t=0.76, p=0.47)$

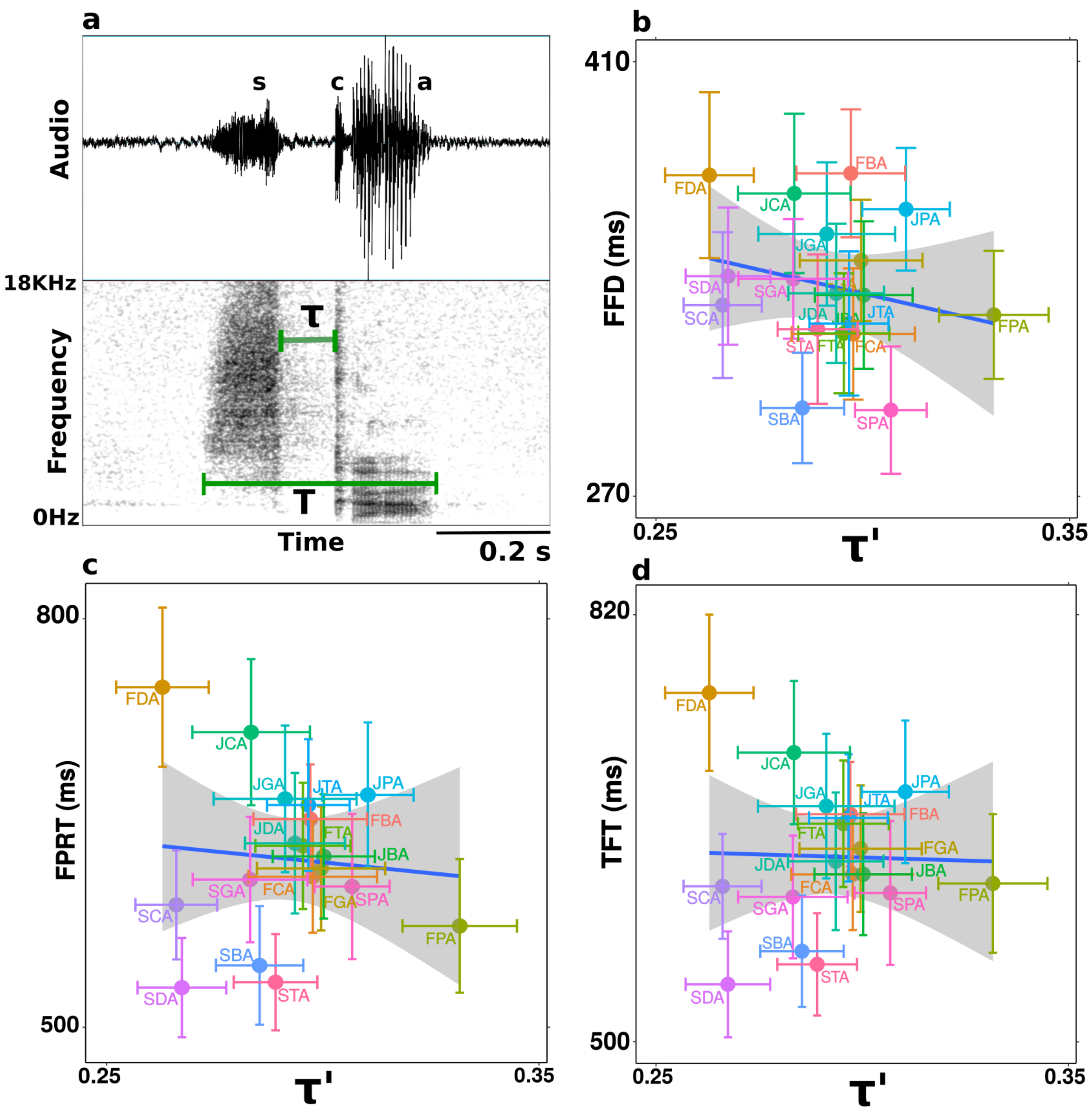


**Figure S2 | Fricative–plosive combinations. a.** Audio and spectrogram example from /SCA/ pronunciation. During these transitions, the vocal tract evolves from a constriction at one point to an occlusion at another one. Transitions are spectrally defined as the interval between the abrupt end of the fricative and the release of the plosive into the vowel [a]. **b - d.** Linear regression test was conducted for the variables TFT, FPRT and FFD with consonantal transition  as the independent variable. Means and standard errors per CCV were considered for the regression as data points come from different block. Any of the variables hold a statistically significant relation with $\tau'$ : FFD ($t=-0.73, p=0.48$) FPRT ($t=0.06, p=0.95$) and TFT ($t=0.27, p=0.79$)
